# The Phenotype-Genotype Reference Map: Improving biobank data science through replication

**DOI:** 10.1101/2022.09.07.506932

**Authors:** Lisa Bastarache, Sarah Delozier, Anita Pandit, Jing He, Adam Lewis, Aubrey C Annis, Jonathon LeFaive, Joshua C. Denny, Robert J. Carroll, Jacob J. Hughey, Matthew Zawistowski, Josh F. Peterson

## Abstract

Population-scale biobanks linked to electronic health record data provide vast opportunity to extend our knowledge of human genetics. While biobanks have already proven their value to research, data quality remains an important concern. Here we introduce the phenotype-genotype reference map (PGRM), a set of 5,879 genetic associations from 523 GWAS publications that can be used for high-throughput replication experiments in biobank data. We tested the PGRM on five ancestry-specific cohorts drawn from four established, independent biobanks and found evidence of robust replications across a wide array of phenotypes. We defined simple replication measures and show how these can be applied to any EHR-linked biobank to detect data corruption and to empirically assess parameters for phenome-wide studies. Finally, we used the PGRM to determine factors associated with reproducibility of GWAS results.

## Introduction

Over the last decade, experimental methods used to study the relationship between genetic variants and human disease at population scale have evolved significantly. Early genome-wide association studies (GWAS) used phenotype-specific cohorts with carefully curated exposures and outcomes.^1,2^ More recent discovery research has gravitated towards multi-purpose biobanks, where phenotypes are defined from a variety of sources, including electronic health records (EHR).^3–7^ As resources with both breadth and depth, biobanks support both GWAS for individual traits and phenome-wide association studies (PheWAS).^8^ “High-throughput” methods, extensive catalogs of GWAS results, and precomputed GWAS x PheWAS associations, have led to a more fine-grained understanding of the genetic underpinnings of complex disease.^9,10^

As the genomics research community moves from using traditional, recruited cohorts towards multi-purpose biobanks, new tools are needed to assess phenotype data quality and ensure replicability of results. High-throughput discovery methods using biobank data are prone to errors related to ascertainment bias, misclassification of phenotypes, errors in sample tracking, and missteps in designing analytic pipelines.^11–13^ While quality control (QC) metrics are routinely applied to the genotype data in biobanks,^14^ analogous metrics are not yet routinely used for phenotypic data.

The abundance of phenotype/genotype associations from prior studies raises the possibility of a new type of QC metric for biobank data, one based on replication. As GWAS results are highly replicable, a new cohort should be able replicate numerous previously discovered associations, given sufficient power.^15,16^ Inability to replicate known associations may indicate data quality issues, problems with phenotype definitions, or other biases that would compromise the generalizability of any new findings from that cohort.^17^ Prior work has demonstrated the feasibility of using replication of known GWAS associations as an indicator of data quality, to validate a new analytic tool, or as a means to compare phenotyping methods.^18–20^

Here, we present the phenotype/genotype reference map (PGRM), a set of genotype associations selected from the National Human Genome Research Institute and European Bioinformatics Institute (NHGRI-EBI) GWAS catalog.^21^ Best practice guidelines for replication studies emphasize the importance of aligning the phenotype definition, cohort composition, and statistical methods used in the initial study with those of the replication study, as well as ensuring adequate power.^22^ To adhere to these recommendations, we manually reviewed all associations and excluded those that were incompatible with a replication study in general purpose biobanks, and included information necessary for power calculations. Phenotypes in the PGRM are represented as phecodes – ICD-based phenotypes that are widely used in biobank studies – to enable automated, phenome-wide replication studies.^23^

To validate the PGRM, we attempted to replicate PGRM associations in four biobanks – BioVU, Michigan Genomics Initiative biobank (MGI), UK Biobank (UKB), and BioBank Japan (BBJ) – finding replication was robust for the three genetic ancestries tested and across disease categories. We developed simplified replication measures that can be applied to any EHR-linked biobank cohort. Through a series of replication experiments, we show how the PGRM can be used to detect data corruption and assess modeling assumptions in phenome-wide studies. Finally, we identified factors that predicted replicability of GWAS associations, including expected factors like reported odds ratio and p-value, and unexpected factors like disease category and the publication date of the original GWAS study. We hope that the PGRM and publicly available source code will enhance the rigor of genetic studies using biobanks.

## Results

### Creating Phenotype-Genotype Reference Map (PGRM)

We curated associations for the PGRM that are compatible with population-scale biobanks, specifically those based on adult populations of both sexes that are not explicitly selected for specific diseases or traits (Figure 1). At the time of download (2022-01-04), the GWAS catalog included 320,477 rows, each specifying an association for a phenotype and genotype. Briefly, we excluded associations based on continuous traits (n=228,853); those with missing statistical information in the catalog (n=63,324); those with a reported P>5×10^−8^ (n=11,805); and associations based on genetic variants that were rare (Minor allele frequency [MAF]<1%) or non-standard genetic variants (e.g. haplotypes) (n=1,530). The remaining 14,897 associations were annotated with 655 experimental factor ontology (EFO) terms – the terminology used by the GWAS catalog to represent diseases and traits – 162 of which had a corresponding phecode (version 1.2) that exactly matched the disease specified in the GWAS catalog. (See Table S1 for EFO→phecode map). Associations that did not have a matching phecode were excluded (n=3,911). We also excluded 2,252 associations that were based on modified phenotypes (e.g., ER-breast cancer; n= 1632), specialized cohorts (e.g., breast cancer in BRCA carriers; n=628) or used non-standard statistical models (n=207).

**Figure 1.**
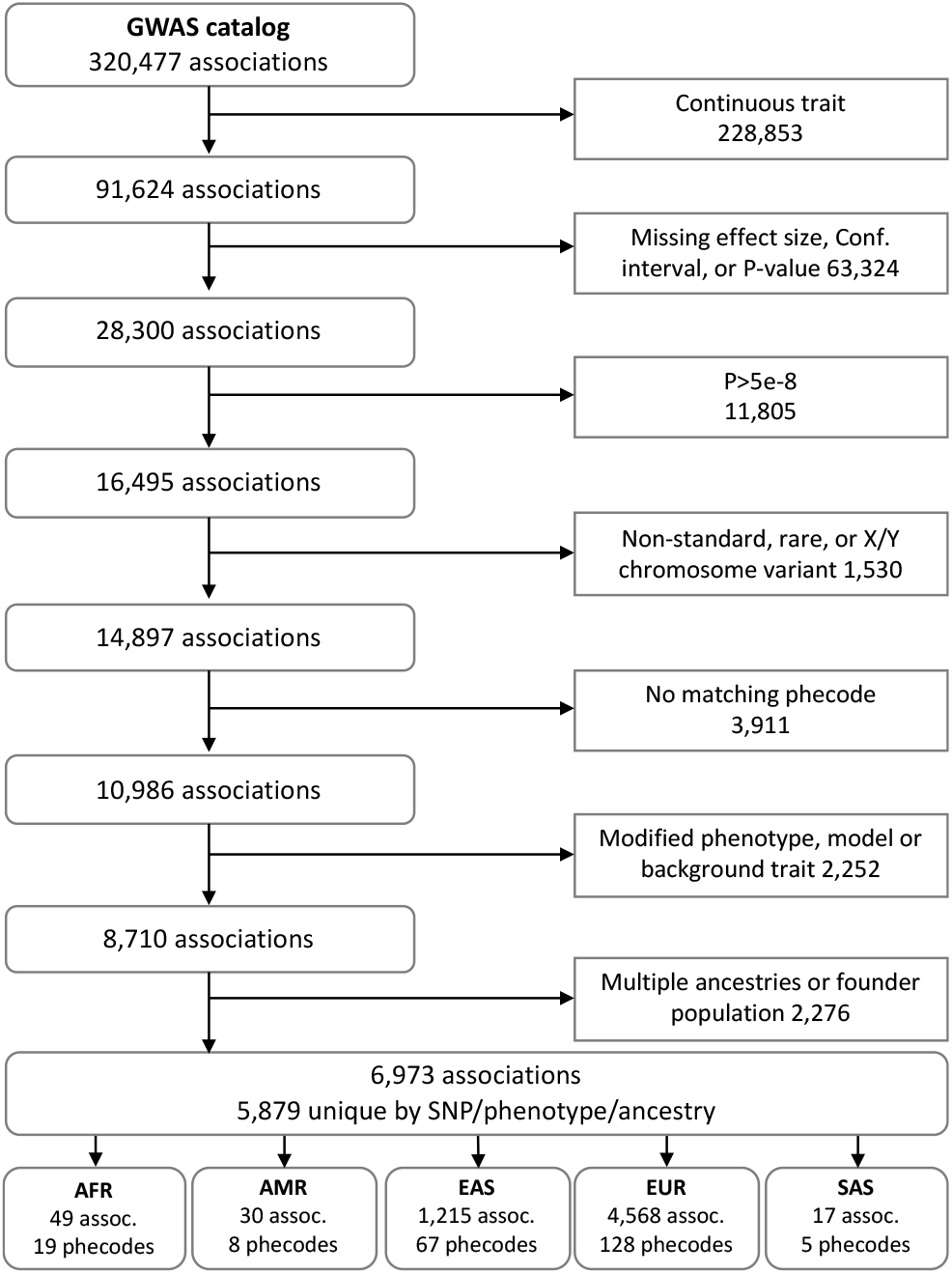
Flow diagram of the creation of the PGRM, beginning with the entire GWAS catalog at the top, to the associations included in the PGRM at the bottom. Rounded boxes on the right show the count of phenotype/genotype associations at each stage, and squares on the left show the number of associations filtered out. The bottom row of the figure shows the number of unique SNP/phenotype associations and unique phecodes included in each subset. Ancestry abbreviations are as follows: AFR=African; AMR= Latino/Admixed American; EAS=East Asian; EUR=European; SAS=South Asian.

Due to differences in allele frequency and linkage disequilibrium patterns between populations, the majority of GWAS findings are ancestry specific.^24^ To facilitate ancestry-match replication studies, we excluded 2,276 associations based on multi-ancestry or founder populations. We annotated PGRM associations according to genetic ancestry of the initial study, including African (AFR; n=49), East Asian (EAS; n=1,215); European (EUR; n=4,568); Latino/Admixed American (AMR; n=30); and South Asian (SAS; n=17). 78% of the associations in the PGRM were based on EUR cohorts.

In total, the PGRM (version 0.0.1) comprises 5,879 unique phenotype/genotype associations drawn from 523 independent GWAS publications (Table S2). These associations capture a wide array of diseases, including 149 unique phecodes from 13 disease categories (Figure S1). The PGRM includes summary statistics from the original study (odds ratios, 95% confidence intervals, and p-values), the PubMed IDs and publication dates, risk alleles, and variant identifiers for builds hg19 and hg38.

### Developing test cohorts of PGRM associations using established biobanks

We aggregated summary statistics from four cohorts: BioVU, MGI, UKB, and BBJ. BioVU was divided into two test cohorts, one of European genetic ancestry (BioVU_Eur_) and another for African genetic ancestry (BioVU_Afr_). (See Table 1 for cohort descriptions and Table S3 for methods used to generate summary statistics).

**Table 1.** Description of biobank test cohorts.

| Test Cohort | Source | Genetic Ancestry | Cohort size | Associations tested | Phenotypes tested |
| --- | --- | --- | --- | --- | --- |
| BioVU<sub>EUR</sub> | BioVU | EUR | 62,777 | 3,268 | 106 |
| MGI | Michigan genomics initiative | EUR | 51,393 | 4,117 | 109 |
| UKB | UK Biobank | EUR | 407,202 | 2,238 | 81 |
| BioVU<sub>AFR</sub> | BioVU | AFR | 12,142 | 31 | 14 |
| BBJ | BioBank Japan | EAS | 178,726 | 384 | 26 |

#### Mapping BBJ phenotypes

The BBJ associations were downloaded from a pre-existing analysis available online (http://pheweb.jp). Unlike the other test cohorts, the BBJ analysis did not use phecodes to define phenotypes. To integrate the BBJ results into the PGRM, we manually mapped 59 of the ICD-based BBJ phenotypes to phecodes so the results could be integrated into the analysis (Table S4).

##### Infobox

**Replication measures & definitions**

###### Replication

An association with a p-value <0.05 & odds ratio in the same direction as the original study.

###### Powered

An association with power ≥ 80% (α=0.05).

###### Overall replication rate (RR_All_)

Number of replicated associations *r*, divided by total number of associations tested *t*.

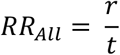

###### Power replication rate (RR_Powered_)

Number of replicated powered associations *r*_*p*_, divided by total number of powered associations tested, *t*_*p*_.

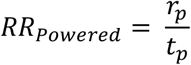

###### Percent powered

Number of powered associations, *t*_*p*_, divided by the number of associations tested, *t*.

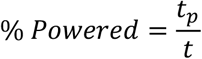

###### Actual over expected ratio (A:E ratio)

Number of replicated associations divided by the sum of the power estimate.

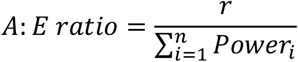

### Developing replication measures for QC and comparative studies

To address the lack of QC tools designed for biobank data, we developed simple replication measures that can be applied to any EHR-biobank. We considered an association replicated if it had p<0.05 and a direction of effect consistent with the original study. Power was defined as having power **≥**80% using the lower 95% confidence interval from the catalog to account for winner’s curse.^25^ The overall replication rate (RR_All_) and powered replication rate (RR_Power_) describe the proportion of PGRM associations replicated by a test cohort. A third metric, the actual:expected ratio (AER), is a similar measure that includes all PGRM associations, regardless of power. This metric may be more suitable for smaller association sets within the PGRM, as is found in under-represented ancestries. (See Infobox for terms and definitions).

#### Excluding associations discovered with test cohorts

We identified 3,180 associations in the PGRM that were discovered using one or more of the test cohorts (Table S5). For these non-independent associations, the overall replication rate (RR_All_) was higher than the RR_All_ of the independent associations (75.1% versus 43.6%) as expected. These non-independent associations were excluded from replication analyses with the test cohorts; over half of the associations derived from BBJ were excluded in this step, reflecting the outsized role this biobank plays in GWAS for the East Asian population.

#### Applying replication measures to test cohorts

The overall replication rate (RR_All_) across test cohorts ranged from 37% - 61%. The replication measures that took power into account were more consistent across cohorts, with the RR_Power_ ranging from 76% - 85% and the AER ranging from 0.78 – 0.94. The AER utilized an average of 3.5 times more associations than the RR_Power_ (Table 2 and Table S6).

**Table 2.** Replication measures for biobank test cohorts. Full results can be found in Table S6.

| Test Cohort | % Power | RR<sub>All</sub> | RR<sub>Power</sub> | A:E ratio |
| --- | --- | --- | --- | --- |
| BioVU<sub>EUR</sub> | 26.0% | 41.4% | 76.3% (651 of 853) | 0.81 |
| MGI | 22.5% | 36.9% | 76.1% (706 of 928) | 0.79 |
| UKB | 38.1% | 56.9% | 85.2% (727 of 853) | 0.94 |
| BioVU<sub>AFR</sub> | 45.2% | 61.3% | 78.6% (11 of 14) | 0.94 |
| BBJ | 57.0% | 53.4% | 76.7% (168 of 219) | 0.78 |

### PGRM demonstrations

We conducted a series of replication experiments to demonstrate applications of the PGRM. In these analyses, we compare the results from the BioVU_Eur_ study (the reference) to results generated with the same dataset after some modification.

#### Detecting data corruption

We hypothesized that corrupted data would negatively impact a cohort’s replication rate, resulting in a lower RR_Power_ and AER. We found that randomizing subjects at 10% increments (starting with a cohort at 0% randomization and ending with a fully randomized cohort) decreased the replication rate monotonically. The RR_All_ and RR_Power_ between 10% increments was statistically significant for all pairs tested. A fully randomized cohort yielded a replication rate of 1.6%, consistent with chance, indicating that the RR_Power_ is sensitive to data corruption. (Figure 2A, Table S7).

**Figure 2.**
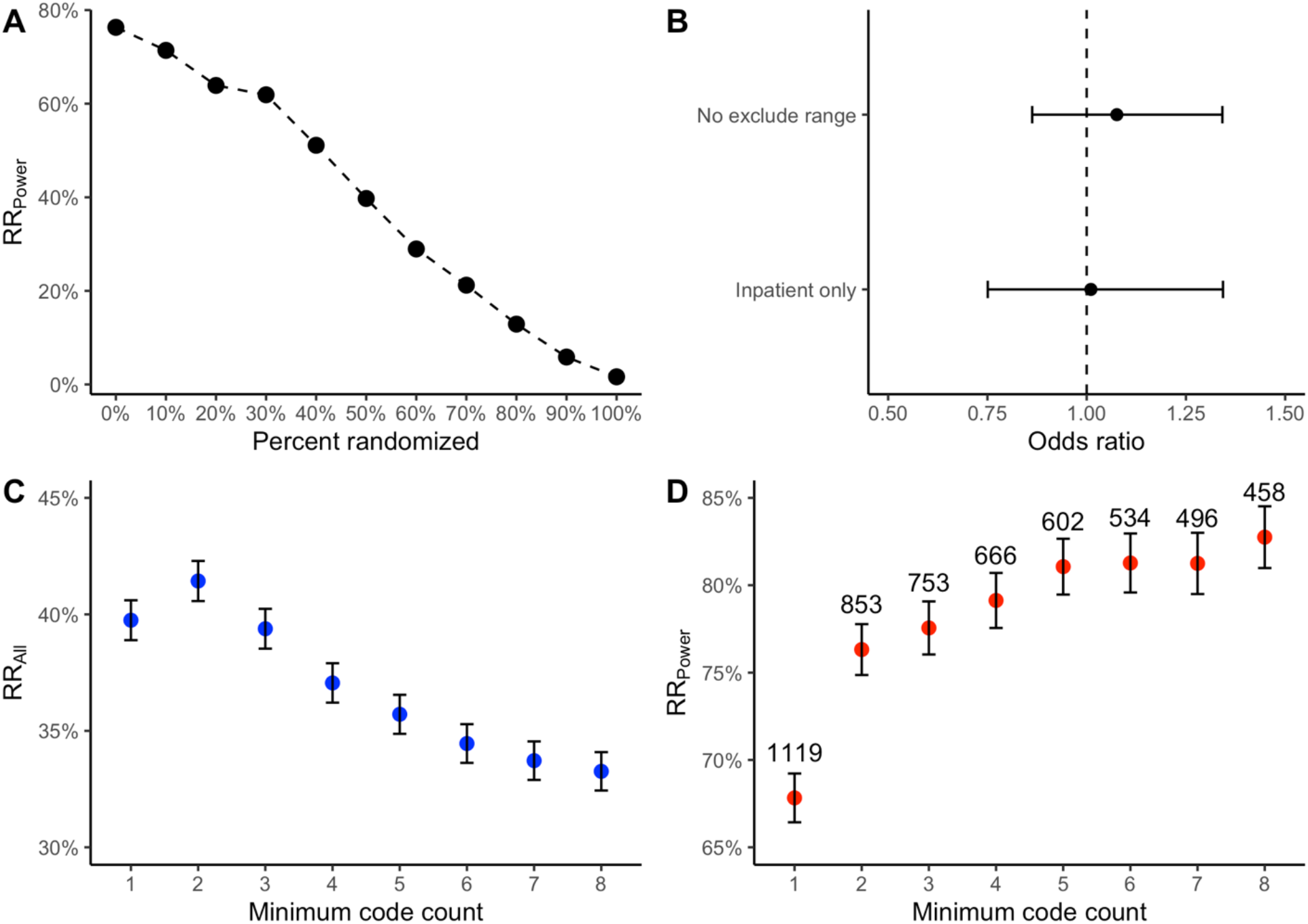
Replication experiments conducted with the PGRM. (A) The replication rate for the BioVU cohort was calculated after permuting subject IDs in the genotype file. The RR_Power_, shown as black circles, decreased significantly with every additional 10% of subjects that were randomized. (B) No significant difference was detected for the odds of replicating in an analysis that did not use exclude ranges (top line) or when only inpatient codes were included (bottom line), when compared to a reference dataset. (C) The RR_All_ for varying minimum code count (MCC) thresholds are shown as blue dots. The maximum RR_All_ was observed with MCC=2. (D) The RR_Power_ for varying MCC, shown as red dots. Above the RR_Power_, the total number of associations in each analysis are labeled. The maximum RR_Power_ was observed for MCC=8; however, this analysis only included 485 associations. The number of powered associations decreased with increasing MCC.

#### Testing utility of exclude ranges

Phenome-wide analyses such as PheWAS typically exclude controls with similar conditions to the target phenotype (e.g. the phecode for “Migraine” excludes controls with “Other headache syndromes”). In principle, excluding controls with similar conditions to the target phenotype should enhance the ability to detect association (by removing potential false negatives), but their effect on replication has never been studied systematically. We hypothesized that the replication would be more robust when using exclude ranges (which were used in the reference analysis) compared to without. We found that replication measures were nominally higher for summary statistics generated with exclude ranges than those without (RR_Power_ = 76.3% versus 75.0%; AER= 0.81 versus 0.79), but this difference was not statistically significant. The number of powered associations and the RR_All_ were also not significantly different (Figure 2B, Table S8A).

#### Assessing the effect of missing phenotype data

Some biobanks only include a subset of ICD codes. For example, until recently the UK biobank included only inpatient ICD codes. While UK biobank has a track record of producing robust results, the effect of missing outpatient codes in a biobank cohort has never formally been assessed. We tested the hypothesis that phenotypes defined using inpatient ICD codes alone (excluding codes from the outpatient context) would decrease both the power and replication rate of a cohort. We found that the using only inpatient codes did significantly reduce the number of powered associations, from 853 to 337, and significantly decreased the RR_All_ (29.5% versus 41.4%; p=4.0×10^−22^). However, the RR_Power_ was not significantly different (Figure 2B, Table S8B).

#### Comparing minimum code count (MCC) thresholds

The default setting in the PheWAS R package requires cases have at least two instances of a phecode, referred to as minimum code count (MCC) of 2. Prior studies found MCC of 2 maximized phenotype accuracy (a balance of precision and recall), but these studies were based on a limited number of phenotypes.^23^ We used the PGRM to assess the effect of different MCC on a phenome-wide scale. Our results showed a tradeoff between power and replication with increasing MCC. The number of powered associations was highest at MCC of 1 (n=1,126) and lowest at MCC=8 (n=458) due to the reduction in cases. The decrease in powered associations was most precipitous from MCC of 1 to 2, where 273 (8%) of associations lost power, and this decrease was statically significant stepwise from MCC of 1 to 4. RR_Power_ increased significantly from MCC of 1 to 2 (67.5% to 76.3%), and continued to increase with ascending MCC, though the stepwise differences were small and not statistically significant. MCC of 2 yielded the most replications overall (n=1,354), suggesting that this threshold strikes a balance between power and phenotype accuracy (Figure 3B; Table S9).

### Assessing replicability with biobank test cohorts

We used our biobank test cohorts to assess the replicability of PGRM associations. First, we looked for associations that were “replication resistant” (i.e. associations that failed to replicate in multiple cohorts). Of the 393 associations that were powered in all three European ancestry cohorts, 22 (5.6%) did not replicate in all three cohorts. Replication resistant associations were not restricted to any one phenotype or phenotype category (See Table S10 for a list of non-replicated associations). 86% of associations (n=339) replicated in at least two cohorts, and 66% (n=258) replicated in all three (Figure S2).

#### Comparing odds ratios

Prior work has shown that GWAS findings are upwardly biased in terms of reported odds ratios (OR).^26^ Therefore, we hypothesized that ORs in the PGRM would be higher than those derived from the test cohort replications. For the 4,373 associations that replicated in the test cohorts, we found that ORs were significantly higher in the PGRM than the test cohort (t-test P=2.1×10^−18^; 95% OR=0.05 [0.039 – 0.061]) (Figure S3).

#### Assessing factors of replication

Using a multivariate logistic regression, we assessed the factors of PGRM associations that were correlated with replication in our test cohorts. As expected, factors related to the strength of the association and power were significantly associated with replication. Higher odds ratio and lower p-values in the original study were positively correlated with replication, as were the number of cases and controls in the test cohort (p<0.001; Table 3).

**Table 3.** Factors associated the replication of association from the GWAS catalog.

| Original association attributes | Odds ratio | 95% CI | P |
| --- | --- | --- | --- |
| Effect size<sup>a</sup> | 1.20 | 1.15 – 1.24 | $5.6 \times 10^{-21}$ |
| P-value<sup>b</sup> | 1.07 | 1.06 – 1.08 | $4.21 \times 10^{-99}$ |
| Risk allele frequency | 1.16 | 0.97 – 1.39 | 0.094 |
| GWAS publication date<sup>c</sup> | 0.92 | 0.90 – 0.93 | $4.8 \times 10^{-28}$ |
| # of times in catalog | 1.17 | 1.06 – 1.29 | $1.7 \times 10^{-3}$ |
| **Test cohort attributes** |  |  |  |
| Cases<sup>d</sup> | 1.14 | 1.12 – 1.16 | $4.8 \times 10^{-54}$ |
| Controls<sup>e</sup> | 1.02 | 1.017 – 1.025 | $7.9 \times 10^{-29}$ |
| **Category** |  |  |  |
| Neoplasms | Reference | - | - |
| Circulatory | 0.86 | 0.72 – 1.04 | 0.118 |
| Dermatologic | 0.60 | 0.49 – 0.73 | $1.0 \times 10^{-6}$ |
| Digestive | 0.64 | 0.55 – 0.75 | $1.4 \times 10^{-8}$ |
| Endocrine | 0.74 | 0.61 – 0.91 | $4.3 \times 10^{-03}$ |
| Genitourinary | 0.78 | 0.53 – 1.14 | 0.199 |
| Infectious disease | 0.44 | 0.24 – 0.84 | 0.012 |
| Metabolic/heme | 1.41 | 0.84 – 2.39 | 0.198 |
| Musculoskeletal | 0.43 | 0.33 – 0.57 | $6.5 \times 10^{-10}$ |
| Neurological | 0.56 | 0.47 – 0.68 | $1.1 \times 10^{-9}$ |
| Psychiatric | 0.12 | 0.10 – 0.16 | $4.9 \times 10^{-60}$ |
| Sense organs | 0.40 | 0.33 – 0.49 | $7.2 \times 10^{-19}$ |

<sup>a</sup> Effect size = log(odds ratio) reported in the catalog. By 0.1 increase
<sup>b</sup> -log<sub>10</sub>(p-value)
<sup>c</sup> In years
<sup>d</sup> Cases in test cohort; Per 1,000 cases
<sup>e</sup> Controls in test cohort; Per 10,000 controls

We found that the number of times an association was reported in the catalog was positively correlated with replication (P=1.7×10^−3^; OR=1.17 [1.06 – 1.29]), as has been previously reported.^18^ The date the association was published was negatively correlated with replication, such that the later the association was first published, the less likely it was to replicate (p=4.8x^-28^; OR=0.92 [0.90 – 0.93] per year, after 2005). Seven disease categories were significantly less likely to replicate when compared with neoplasm phenotypes (the largest category of association and the category with the highest replication rate), including digestive, endocrine, genitourinary, metabolic, psychiatric, and sense organs. Psychiatric disorders had a notably low likelihood of replicating (p=5×10^−60^; OR=0.12 [0.10 – 0.16]).

## Discussion

The PGRM is a large set of phenotype/genotype associations that can be used to conduct replication studies at a phenome-wide scale. Phenotypes in the PGRM were annotated with phecodes to ensure interoperability with EHR-linked biobanks, and associations were curated for compatibility with general-purpose biobank cohorts. Each PGRM association includes information about the ancestry of the original cohort and summary statistics to facilitate power calculations. In total, the PGRM includes 5,879 replication candidates (unique by SNP, phenotype, and ancestry) drawn from 523 GWAS studies, and spanning 149 distinct diseases from across the medical phenome.

Given the high reproducibility of GWAS findings, we anticipated that PGRM associations would robustly replicate in our five mature biobank cohorts. Indeed, the overall phenome-wide replication rates were higher than previously reported, likely due to the larger cohorts tested in this study.^18^ The majority (86%) of phenotypes replicated at least once, indicating the inclusion of many phenotypes adds robustness. Phenome-wide replications have not been studied across genetic ancestry cohorts. Encouragingly, we found replication rates were comparable across ancestry groups tested. The powered replication rate was strikingly consistent across tests cohorts, averaging 79% across all cohorts with a range from 76% - 85%. These results provide a benchmark against which new test cohorts can be compared.

Through a series of demonstration studies, we showed how the PGRM can be used to assess data quality and analytical assumptions. We found that replication measures are sensitive to data corruption, indicating that they may be used to detect data problems that commonly occur in the transfer and processing of large datasets. We also showed how the PGRM can be used to test the effect of different analytical parameters used in PheWAS, such as exclude ranges and minimum code count threshold, and assess the effect of missing phenotype data.

The PGRM offers a way for researchers to assess phenome-wide parameters in their own datasets. This is important because the optimal settings for a phenome-wide analysis will differ depending on the goals of the analysis and the characteristics of the underlying dataset. Some analyses might benefit from enhanced precision, and others from increased power. Furthermore, biobank cohorts (and sub-cohorts therein) differ in terms of longitudinally and density, which means that the effect of various parameters like minimum code count will likely differ as well. Therefore, the optimal parameters for an analysis are study dependent. While replication measures do not tell the whole story, the PGRM is at least one way of generating empirical evidence for decisions analysts must make when modeling phenome-wide data.

When interpreting PGRM measures, investigators should be aware that there are multiple reasons beyond data quality for why a test cohort may yield a low replication rate. Ideally, a replication is attempted using the same phenotype and a population that is comparable in terms of genetic background, environmental exposures, and demographics.^22^ But there is much complexity to unpack in this seemingly simple statement. What makes two cohorts equivalent? How might we ensure that two phenotypes are indeed the same? Through a manual effort, we attempted to align the PGRM associations with the characteristics of a general-purpose biobank. However, because there is no consistent framework used to specify these attributes, important details may be missing in the GWAS catalog. Moreover, the relationship between genotype, environment, and phenotype is so complex that the key conditions necessary for replication are not always known. Future work may use replication experiments to further elucidate how differences in phenotype definition and cohort compositive affect replicability and generalizability.

In addition to supporting quality measures, the PGRM enabled us to study the GWAS catalog itself, identifying factors that influence the replicability of GWAS findings. Unsurprisingly, we found factors relating to power strongly influenced replicability. While replication occurred across all 13 disease categories tested, not all disease categories were equally likely to replicate. Psychiatric phenotypes were a outlier in this analysis, with a 0.12 odds of replicating relative to neoplasm phenotypes. This strikingly low replication rate, which was observed across our test cohorts, may be due to challenges capturing psychiatric phenotypes at scale using EHR data.^27,28^ We also found that associations from more recent publications were less likely to replicate, with a 0.90 odds of replication per year since 2005. The reasons for the decreased replication rate over time may be partially explained by the early discovery of the most robust GWAS associations. But it also may be linked to recent concerns that decreased transparency, methodological rigor, and the use of non-representative cohorts is eroding GWAS’s impressive track record of replicability.^29,30^

The PGRM is not without limitations. First, non-European ancestry associations are very much under-represented in the catalog, a reflection of the European ancestry bias of GWAS studies.^31^ We hope that calls to increase diversity in genetic research will lead to a more complete picture of genetic associations in under-studied ancestries.^32,33^ The PGRM is designed to expand as additional diverse ancestry associations are discovered. Furthermore, recent advances in trans-ancestry GWAS methodology may be used to generate ancestry-agnostic entries in the PGRM.^34^

Second, the PGRM is limited to phecode-based phenotypes. Thus, continuous traits, which make up the majority of the GWAS catalog, are not currently included. Future iterations could integrate measurements such as height and laboratory results to facilitate replication experiments on continuous traits.^35^ The PGRM could also evolve to use data sources beyond ICD codes, such as survey responses. Each addition will require a new map, like the one we created for BBJ phenotypes in this analysis, but existing features will still be applicable.

Replication is a powerful way to assess the quality of large, complex biobank datasets, and to better understand analytical assumptions used in modeling phenome-wide data. To facilitate the use of the PGRM, we created a publicly available R package that allows the PGRM to be incorporated into existing biobank QC pipelines or used to conduct replication experiments. We hope that the development of this resource can help maintain the high standards of reproducibility in the biobank era.

## Methods

Files from the publicly available GWAS catalog were downloaded on January 4, 2022 (“All associations v1.0.2,” “All studies v1.0.2,” and “All ancestry data v1.0”). Each row of the “All association” file represents a genotype/phenotype association. We included SNP/phenotype associations that met the following criteria: (1) were based on a single nucleotide polymorphism (SNP) (as opposed to a haplotype or combination of SNPs), excluding the X or Y chromosome; (2) had a specified odds ratio, and confidence intervals; (3) reported a p-value of < 5e-8; and (4) were based on a binary trait modeled as a logistic regression. We excluded continuous traits like blood pressure by using the OR_or_BETA column (all values < 1), mentions of measurement words (e.g “increase”, “decrease”, “ratio”) in the 95_CI_TEXT, and specification of continuous trait in the DISEASE_TRAIT or P-VALUE (TEXT) field (e.g. “age of onset”).

### Mapping phenotypes

The GWAS catalog diseases and traits are annotated with the Experimental Factor Ontology (EFO).^36^ We attempted to annotate all EFO terms present in the filtered list of associations with a matching phecode. We used WikiMedMap to generate candidate matches, and manually chose matching phenotypes.^37^ The full map can be found in Table S1.

### Alternative models, background traits, and modified phenotypes

Although the conditions necessary to ensure replication across cohorts are not well characterized, prior work has indicated that misalignment of study design, cohort composition, and phenotype definition decreases the likelihood of replication. Thus, to maximize the likelihood of replication in a test cohort, we filtered out GWAS catalog associations with characteristics that were not aligned with a general population biobank cohort. We excluded associations from the PGRM for the following reasons:

1. Phenotype misalignment: The catalog includes phenotypes that are qualified by severity (e.g. mild proteinuria), family history (e.g. familial lung cancer), age of onset (e.g. Early-onset schizophrenia), and subtype (e.g. ER-breast cancer).
2. Cohort misalignment: Some catalog associations are based on specialized cohorts wherein all study participants share a characteristic, defined on the basis of disease (e.g. diabetic nephropathy in type 1 diabetes), exposure (e.g. Clostridium difficile infection in antibiotics-users), genetics (e.g. breast cancer in individuals with pathogenic BRCA1 variants), sex, (e.g. hypertension in males), and young age (e.g. obesity in children). These are referred to as “background traits” in the GWAS catalog, and they are sometimes (though not always) annotated in the column of that name.
3. Non-standard statistical models: While GWAS results are typically based on logistic regression and additive genotype, the catalog includes recessive or dominant associations models as well those generated from non-standard models (e.g. family based models, interactions).

We manually extracted and normalized information regarding the phenotype, study cohort, and statistical model using information from the following columns in the GWAS catalog: DISEASE_TRAIT, P-VALUE (TEXT), STUDY, and BACKGROUND_TRAIT, as well as the study title. When necessary, the original manuscript was reviewed to resolve ambiguous information provided in these columns.

### Ancestry annotation

We annotated study cohort ancestry using the INITIAL SAMPLE SIZE and REPLICATION SAMPLE SIZE in the “All ancestry data” file, as well as the P-VALUE (TEXT) column. We used the 1000 Genomes superpopulations ancestry groupings – African (AFR), East Asian (EAS), European (EUR), AdMixed American (AMR), South Asian (SAS) – as well as Multiple (for cohorts that included >1 genetic ancestry superpopulation), and Other (including individuals from founder populations or genetic ancestries not covered in the superpopulations). We excluded catalog associations that were based on multiple ancestries or included founder populations.

### Consolidating the PGRM

Each association in the PGRM is unique by phenotype, SNP, and ancestry. When associations were reported multiple times in the catalog, we chose the association with the lowest p-value for inclusion in the consolidated PGRM. The number of times an association appeared in the catalog was stored.

### Variant mapping

We used the Ensembl REST API (https://rest.ensembl.org/) to annotate each rsID present in the filtered list of associations. We recorded the allele_string, the location (chromosome and start/end position) and reference allele frequencies from gnomAD (EUR, AFR, EAS, SAS, AMR). For each variant, we stored the unique alleles defined in gnomAD and 1000 genomes. A variant ID was created for each non-multi-allelic variant by concatenating the chromosome, start position, reference allele and alternate allele (e.g. 1:62782860:T:C). Each row in the catalog was annotated with a variant ID. Variants with more than two alleles specified were flagged as “multi-allelic” and excluded if the risk allele was ambiguous in the catalog. The annotation process was conducted twice: first with build GRCh37 (http://grch37.rest.ensembl.org) and then for build GRCh38 (http://rest.ensembl.org).

### Risk allele

The risk allele is defined as the allele that is associated with risk of the phenotype (i.e. odds ratio > 1). For each association, we annotated the risk allele as reference (“ref”) or alternative (“alt”) according to the STRONGEST_SNP_RISK_ALLELE reported in the catalog. In cases where the catalog did not report a STRONGEST_SNP_RISK_ALLELE, we determined the risk allele by matching allele frequencies in gnomAD with the risk allele frequency (RAF) reported in the catalog, matching by ancestry. When the RAF and risk allele were absent from the catalog or ambiguous, we labeled the direction as “unknown.” Following association testing, we check the allele direction against the results of our five test cohorts. For associations with an unknown direction, if two or more test cohorts replicated the association at p<0.05 with the same direction of effect, or if a single cohort replicated an association with p<0.01, we set the PGRM direction accordingly. Some directions were changed based on rest results. Associations without a clear risk allele were labeled with a “?” and treated as “alternate” in subsequent calculations. Each row was annotated with the source of information on the allele direction (e.g. Risk allele, RAF, one or more test cohorts, or unknown).

### Source cohort annotation

We identified all associations that subjects from BioVU (or eMERGE), MGI, or UK biobank by systematically searching for the biobank names in the source manuscripts. This annotation was used in subsequent replication measures to prevent testing for self-replication (i.e. replicating an association from UK biobank with the UK biobank cohort).

### Phenotype Category labels

Each phenotype in the PGRM is labeled by one of 13 category labels. These labels were derived from the phecode category labels, which were modified for the purposes of this study. The metabolic/endocrine categories split into two separate categories. Because there were very few hematopoietic phenotypes, these were consolidated into the metabolic category. The “mental disorders” category was renamed “psychiatric” and phenotypes for Alzheimer’s disease and dementia were to the “neurological” category.

## Generating test cohort results

We derived summary statistics for associations in the PGRM in four mature biobanks: BioVU, MGI, UKB, and BBJ. The size, ancestry backgrounds, and descriptions of each cohort and phenotype definitions are listed in Table S3 and details on the methods used to calculate associations are described for each biobank in the following paragraphs. A complete set of association results with annotations can be found in table S6.

### BioVU

The BioVU cohort comprised adults (age >=18 at last visit) who were genotyped on the MEGA array. Variants were imputed using the 1000genomes. Imputed variants were converted into hard calls, excluding those with a R2 < 0.85.^38^ Using ancestry informative markers, we defined a cohort of African genetic ancestry and another of European genetic ancestry (BioVU_AFR_ and BioVU_EUR_). From these two cohorts, we excluded SNPs with minor allele frequency of <1% and those with missingness > 2% or a Hardy-Weinberg p-value < 1e-5 as described by Coleman et al.^39^ Phenotypes were defined using phecodes Version 1.2.^40^ Cases were defined as having at least two phecodes on unique dates (referred to as minimum code count [MCC] = 2), and controls were defined using exclude ranges.^23^ We used the run_PGRM_assoc() function from the pgrm R package to conduct association tests for each phenotype/SNP pair in the PGRM, adjusting for age, sex, and the first eight genetic principle components (PCs). The run_PGRM_assoc() using logistic regression to conduct the association tests with raw genotype and phenotype data for a test cohort. This analysis will be referred to as the benchmark results in subsequent studies.

### MGI cohort

The Michigan Genomics Initiative (MGI) has previously been described.^41^ Briefly, MGI is a biobank of Michigan Medicine patients with linked electronic health records and genetic data. Participants were genotyped using an Illumina Infinium CoreExome-24 bead array platform and imputed using the TOPMed haplotype reference panel. Binary phecode phenotypes were created based on ICD inclusion and exclusion criteria using the PheWAS R package v0.99.5.-5.^42^ We used the default PheWAS package parameters of MCC of 2 and exclude ranges. We performed a GWAS in a set of 51,583 MGI samples of genetically inferred European ancestry on a total of 1,712 PheCode traits with case count ≥20. GWAS are run using a mixed model implemented in SAIGE v0.43.3 to account for relatedness and case-control imbalance (Zhou et al., 2020). Variants with R2<0.3 or MAF < 0.01% were excluded. We adjusted for age, inferred sex, genotyping array, and the first ten genetic PCs for each association test. We extracted associations present in the PGRM from the summary statistics of this GWAS x PheWAS analysis.

### UK Biobank

We performed GWAS on a set of “White British” participants drawn from the full UK Biobank (UKB) cohort released in July 2017. These results were generated under UK Biobank data application number 24460. UKB participants are genotyped on an Affymetrix Axiom array, then imputed using the TOPMed imputation reference panel. A total of ∼37,000,000 high quality variants remained for GWAS analysis after removing variants with R2≤0.3 and/or MAF≤0.01%.^43–45^ We identified individuals of “White British” ancestry using the original definitions provided by UKB. We computed phecodes using on including diagnosis codes using the PheWAS R package^42^ and collected covariates sex, genotyping batch, and age from the participant electronic health records. We used 1,418 of the resulting phenotypes having case count >=100 in the analysis in a total of 407,202 participants. We tested for association between genetic markers and case-control status for each phecode trait using a logistic mixed model implemented in the SAIGE association software.^46^ Covariates included age, sex, chip version, and the first four principal components. We extracted associations present in the PGRM from the summary statistics of this GWAS x PheWAS analysis.

### Biobank Japan (BBJ) cohort

We accessed the summary statistics of the BBJ results from the website https://pheweb.jp/. The website hosts GWAS results from 229 phenotypes, 42 of which are based on ICD-10 codes and described in a publication by Sakaue et. al.^9^ We manually identified 59 phenotypes in the BBJ that were exact matches to phecodes in the PGRM. (Table S4). We downloaded the summary statistics for each phenotype, then extracted summary statistics for the SNP/phenotype associations in the PGRM, including the beta, standard error, and p value.

### Annotating test cohorts

We annotated the summary statistics of each test cohort using the annotate_results() function from the pgrm R package. This function merges association results from a test cohort with the PGRM, filtering by the specified ancestry, and annotates the result set with the following information:

1. Information from the original GWAS study, including the accession number, summary statistics, risk allele direction.
2. Power for each association based on the case/control counts from the test cohort, the ancestry-matched allele frequency from gnomAD, and the lower confidence interval from the GWAS catalog. Power is calculated for alpha=0.05, and
3. Replication Boolean value, set to 1 if the test cohort replicated the association. Replication is defined as having P < 0.05 and odds ratio in the same direction as reported in the GWAS catalog.
4. The comparison of the confidence intervals (CIs) from the original study and test cohort. Associations are labeled “overlap” if the CIs from catalog and test association are overlapping, “test_cohort_greater” if the lower CI from the test cohort is higher than the upper CI of the catalog cohort, and “PGRM_greater” if the lower CI from the PGRM is higher than the upper CI of the test cohort.

### Power calculations

All power calculations were performed using the genpwr.calc function in the genpwr R package with an alpha=0.05. We used the lower confidence from the PGRM and the allele frequency (AF) from gnomAD. The lower confidence interval was used instead of the point estimate to compensate for the “winner’s curse”.

### Calculating replication measures in test cohorts

We calculated overall replication rate (RR_All_) and powered replication rate (RR_Power_) by applying the get_RR() function to the annotated test cohorts. We calculated the Actual:Expected ratio (AER) by applying the get_AER() function to each test cohort. The AER is defined as the total number of replicated associations divided by the sum of the power over all associations, a measure used by Palmer and Pe’er.^26^

## Replication experiments

We conducted replication experiments by creating new datasets with modified phenotype files using the BioVU_EUR_ test cohort. We compared these modified datasets to the benchmark dataset to assess the effects of various phenotype and cohort definitions on replication measures. The following list describes the way we modified the BioVU_EUR_ cohort to generate the test datasets.

1. *Randomized phenotypes*: We created randomized cohorts using the BioVU_EUR_ cohort by randomly shuffling a proportion of the individuals in the phenotype and covariate file, but not the genotype file. Shuffling was accomplished with the sample() function in the R base package, without replacement. We created 9 additional cohorts, randomizing from 10% to 100%, at 10% intervals.
2. *No exclude ranges*: We generated a phenotype file without exclude ranges. The benchmark analysis used exclude ranges.
3. *Inpatient only*: We generated a phenotype file using only ICD codes from the inpatient context. The benchmark analysis used ICD codes from all clinical context (inpatient, outpatient, emergency)
4. *Variable minimum code counts:* We defined seven additional phenotype files using different minimum code counts (MCC). The MCC is the number of unique dates a phecode in an individual’s record for them to be classified as cases. The benchmark analysis used MCC=2, and we analyzed phenotype files defined with MCC=1 and 3-8.

Summary statistics were generated for each new dataset and annotated using the run_PGRM_assoc() and annotate_results() functions from the pgrm package. We compared the results of the test datasets against the benchmark BioVU_EUR_ results using the compare_results_sets() function in the pgrm R package. This function assesses the difference between replication measures of two result sets. A Fisher’s exact test compares the results of a benchmark and modified cohort for each measure (RR_All_, RR_Power_, and % Power). The full results of the randomization study can be found in table S7. The results for the exclude ranges and inpatient studies are in table S8, and the results for the MCC study are in table S9.

### Assessing the contents of the PGRM with test cohorts

#### Comparing replications across test cohorts

We assessed the overlap of successful replications across the three European test cohorts, including only associations that were included in all three cohorts, and identified associations that were not replicated in any of the three cohorts. Associations that failed replication across the three cohorts can be found in table S10.

#### Comparing odds ratios of initial versus replication studies

We compared the odds ratios in the PGRM with those generated in our test cohorts using a paired t-test. Only associations that replicated in the test cohorts were included in the analysis.

#### Factors of replication

We used a logistic regression model to assess the factors associate with replication across all five test cohorts, including the following independent variables in the model: risk allele frequency, number of cases from the test cohort, lower CI from the catalog, first year the association was published, the number of times the association occurred in the catalog, the phenotype category, and the ancestry. The full model results are included in table 2.

## Data availability

The PGRM is available in supplementary table 2. Summary statistics from the five test cohorts are available in supplementary table S6.

## Code availability

The PGRM is available as an R package at https://github.com/hugheylab/pgrm. The package includes functions to calculate replication measures from raw datafiles or summary data. All results from test cohorts are included in the R package, to facilitate comparative studies.

## Acknowledgements

This work was supported by the National Library of Medicine grant R01-LM010685 and National Center for Advancing Translational Sciences grant 5UL1TR002243-03. The authors acknowledge the Michigan Genomics Initiative participants, Precision Health at the University of Michigan, the University of Michigan Medical School Central Biorepository, and the University of Michigan Advanced Genomics Core for providing data and specimen storage, management, processing, and distribution services, and the Center for Statistical Genetics in the Department of Biostatistics at the School of Public Health for genotype data curation, imputation, and management in support of the research reported in this publication.

## Supplementary information

Table S1 – EFO to phecode map

Table S2 – PGRM

Table S3 – Summary of test cohorts

Table S4 – BBJ to phecode map

Table S5 – PGRM associations that used test cohort in original study

Table S6 – All summary statistics for test cohorts

Table S7 – Results of randomization experiment

Table S8 – Results of exclude range and inpatient only experiment

Table S9 – Results of minimum code count experiment

Table S10 – Replication resistant associations

Figure S1 – Associations in the PGRM, by phenotype and ancestry

Figure S2 – Overlap of associations between test cohorts

Figure S3 – Odds ratio comparison between original study and replication

## Bibliography

1. Uffelmann, E. et al. Genome-wide association studies. Nat Rev Methods Primers 1, 59 (2021).

2. The Wellcome Trust Case Control Consortium. Genome-wide association study of 14,000 cases of seven common diseases and 3,000 shared controls. Nature 447, 661–678 (2007).

3. Gaziano, J. M. et al. Million Veteran Program: A mega-biobank to study genetic influences on health and disease. J Clin Epidemiol 70, 214–223 (2016).

4. All of Us Research Program Investigators et al. The ‘All of Us’ Research Program. N Engl J Med 381, 668–676 (2019).

5. Leitsalu, L. et al. Cohort Profile: Estonian Biobank of the Estonian Genome Center, University of Tartu. Int J Epidemiol 44, 1137–1147 (2015).

6. Sudlow, C. et al. UK Biobank: An Open Access Resource for Identifying the Causes of a Wide Range of Complex Diseases of Middle and Old Age. PLOS Medicine 12, e1001779 (2015).

7. Nagai, A. et al. Overview of the BioBank Japan Project: Study design and profile. J Epidemiol 27, S2–S8 (2017).

8. Bastarache, L., Denny, J. C. & Roden, D. M. Phenome-Wide Association Studies. JAMA 327, 75–76 (2022).

9. Sakaue, S. et al. A cross-population atlas of genetic associations for 220 human phenotypes. Nat Genet 53, 1415–1424 (2021).

10. Neale lab - aUK Biobank. Neale lab http://www.nealelab.is/uk-biobank.

11. Zuvich, R. L. et al. Pitfalls of merging GWAS data: lessons learned in the eMERGE network and quality control procedures to maintain high data quality: Pitfalls of Merging GWAS Data: Lessons Learned. Genet. Epidemiol. 35, 887–898 (2011).

12. Colhoun, H. M., McKeigue, P. M. & Smith, G. D. Problems of reporting genetic associations with complex outcomes. The Lancet 361, 865–872 (2003).

13. DeLozier, S. et al. Phenotyping coronavirus disease 2019 during a global health pandemic: Lessons learned from the characterization of an early cohort. Journal of Biomedical Informatics 117, 103777 (2021).

14. Purcell, S. et al. PLINK: a tool set for whole-genome association and population-based linkage analyses. Am. J. Hum. Genet. 81, 559–575 (2007).

15. O’Sullivan, J. W. & Ioannidis, J. P. A. Reproducibility in the UK biobank of genome-wide significant signals discovered in earlier genome-wide association studies. Sci Rep 11, 18625 (2021).

16. Marigorta, U. M., Rodríguez, J. A., Gibson, G. & Navarro, A. Replicability and Prediction: Lessons and Challenges from GWAS. Trends Genet 34, 504–517 (2018).

17. Huffman, J. E. Examining the current standards for genetic discovery and replication in the era of mega-biobanks. Nat Commun 9, 5054 (2018).

18. Denny, J. C. et al. Systematic comparison of phenome-wide association study of electronic medical record data and genome-wide association study data. Nat. Biotechnol. 31, 1102–1110 (2013).

19. Denaxas, S. et al. UK phenomics platform for developing and validating electronic health record phenotypes: CALIBER. J Am Med Inform Assoc 26, 1545–1559 (2019).

20. Song, W., Huang, H., Zhang, C.-Z., Bates, D. W. & Wright, A. Using whole genome scores to compare three clinical phenotyping methods in complex diseases. Scientific Reports 8, 11360 (2018).

21. Buniello, A. et al. The NHGRI-EBI GWAS Catalog of published genome-wide association studies, targeted arrays and summary statistics 2019. Nucleic Acids Res 47, D1005–D1012 (2019).

22. NCI-NHGRI Working Group on Replication in Association Studies et al. Replicating genotype-phenotype associations. Nature 447, 655–660 (2007).

23. Bastarache, L. Using Phecodes for Research with the Electronic Health Record: From PheWAS to PheRS. Annual Review of Biomedical Data Science 4, 1–19 (2021).

24. Martin, A. R. et al. Human Demographic History Impacts Genetic Risk Prediction across Diverse Populations. Am J Hum Genet 100, 635–649 (2017).

25. Xiao, R. & Boehnke, M. Quantifying and correcting for the winner’s curse in quantitative-trait association studies. Genetic Epidemiology 35, 133–138 (2011).

26. Palmer, C. & Pe’er, I. Statistical correction of the Winner’s Curse explains replication variability in quantitative trait genome-wide association studies. PLoS Genet 13, e1006916 (2017).

27. Curtis, D. Analysis of 50,000 exome-sequenced UK Biobank subjects fails to identify genes influencing probability of developing a mood disorder resulting in psychiatric referral. J Affect Disord 281, 216–219 (2021).

28. Li, Z. et al. Validation of UK Biobank data for mental health outcomes: A pilot study using secondary care electronic health records. International Journal of Medical Informatics 160, 104704 (2022).

29. Burt, C. & Munafò, M. Has GWAS lost its status as a paragon of open science? PLOS Biology 19, e3001242 (2021).

30. Munafò, M. R., Tilling, K., Taylor, A. E., Evans, D. M. & Davey Smith, G. Collider scope: when selection bias can substantially influence observed associations. Int J Epidemiol 47, 226–235 (2018).

31. Morales, J. et al. A standardized framework for representation of ancestry data in genomics studies, with application to the NHGRI-EBI GWAS Catalog. Genome Biol 19, 21 (2018).

32. Manolio, T. A. Using the Data We Have: Improving Diversity in Genomic Research. Am J Hum Genet 105, 233–236 (2019).

33. Fatumo, S. et al. A roadmap to increase diversity in genomic studies. Nat Med 28, 243–250 (2022).

34. Peterson, R. E. et al. Genome-wide Association Studies in Ancestrally Diverse Populations: Opportunities, Methods, Pitfalls, and Recommendations. Cell 179, 589–603 (2019).

35. Goldstein, J. A. et al. LabWAS: Novel findings and study design recommendations from a meta-analysis of clinical labs in two independent biobanks. PLoS Genet 16, e1009077 (2020).

36. Malone, J. et al. Modeling sample variables with an Experimental Factor Ontology. Bioinformatics 26, 1112–1118 (2010).

37. Sulieman, L., Wu, P., Denny, J. & Bastarache, L. WikiMedMap: Expanding the Phenotyping Mapping Toolbox Using Wikipedia. bioRxiv 727792 (2019) doi:10.1101/727792.

38. Johnson, E. O. et al. Imputation across genotyping arrays for genome-wide association studies: assessment of bias and a correction strategy. Hum Genet 132, 509–522 (2013).

39. Coleman, J. R. I. et al. Quality control, imputation and analysis of genome-wide genotyping data from the Illumina HumanCoreExome microarray. Brief Funct Genomics 15, 298–304 (2016).

40. https://phewascatalog.org/.

41. Zawistowski, M. et al. The Michigan Genomics Initiative: a biobank linking genotypes and electronic clinical records in Michigan Medicine patients. 2021.12.15.21267864 Preprint at https://doi.org/10.1101/2021.12.15.21267864 (2021).

42. Carroll, R. J., Bastarache, L. & Denny, J. C. R PheWAS: data analysis and plotting tools for phenome-wide association studies in the R environment. Bioinformatics 30, 2375–2376 (2014).

43. McCarthy, S. et al. A reference panel of 64,976 haplotypes for genotype imputation. Nat Genet 48, 1279–1283 (2016).

44. Taliun, D. et al. Sequencing of 53,831 diverse genomes from the NHLBI TOPMed Program. Nature 590, 290–299 (2021).

45. Kowalski, M. H. et al. Use of >100,000 NHLBI Trans-Omics for Precision Medicine (TOPMed) Consortium whole genome sequences improves imputation quality and detection of rare variant associations in admixed African and Hispanic/Latino populations. PLoS Genet 15, e1008500 (2019).

46. Zhou, W. et al. Efficiently controlling for case-control imbalance and sample relatedness in large-scale genetic association studies. Nat Genet 50, 1335–1341 (2018).

