## Supplemental figures 1-3 for "The Phenotype-Genotype Reference Map: Improving biobank data science through replication"

**Figure S1A**. There are 28 phenotypes that occur in the PGRM at least 50 times. These phenotypes are show here, sorted by the number of SNPs in the PGRM. The color of the bars indicates the genetic ancestry of the association.

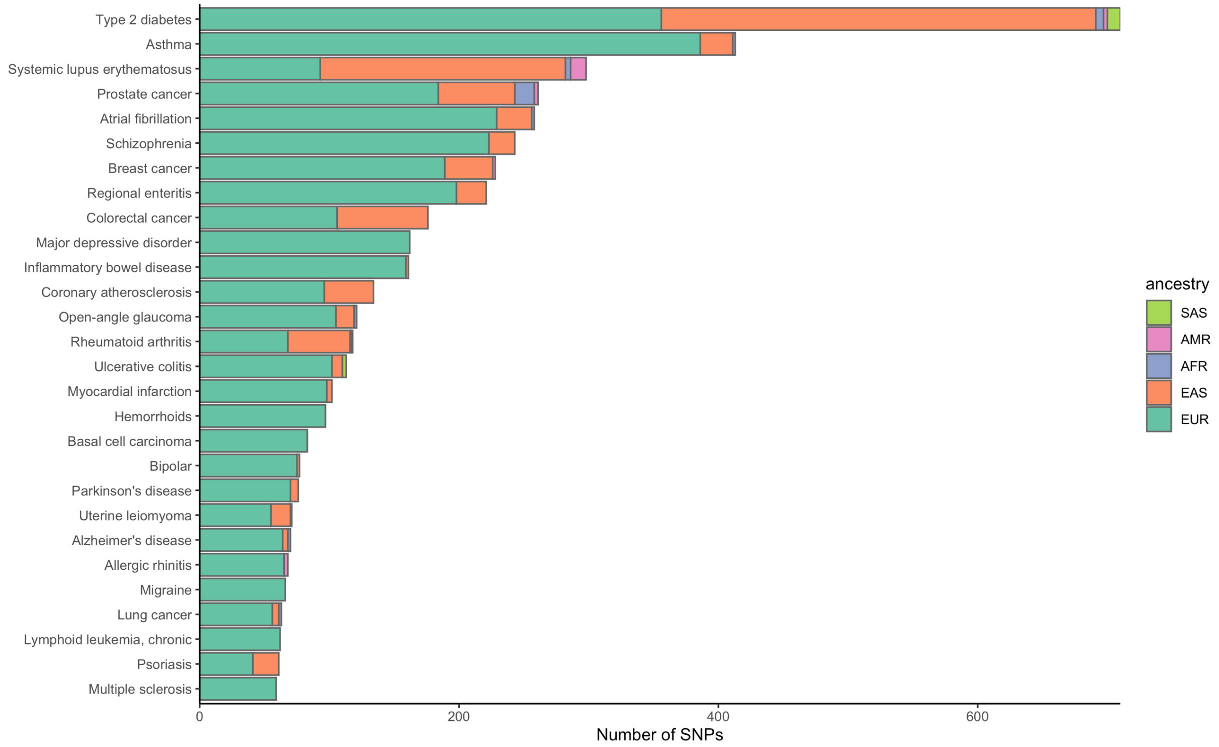

**Figure S1B**. Phenotypes in the PGRM with fewer than 50 SNPs, organized by phecode category and sorted by the number of times they appear in the PGRM.

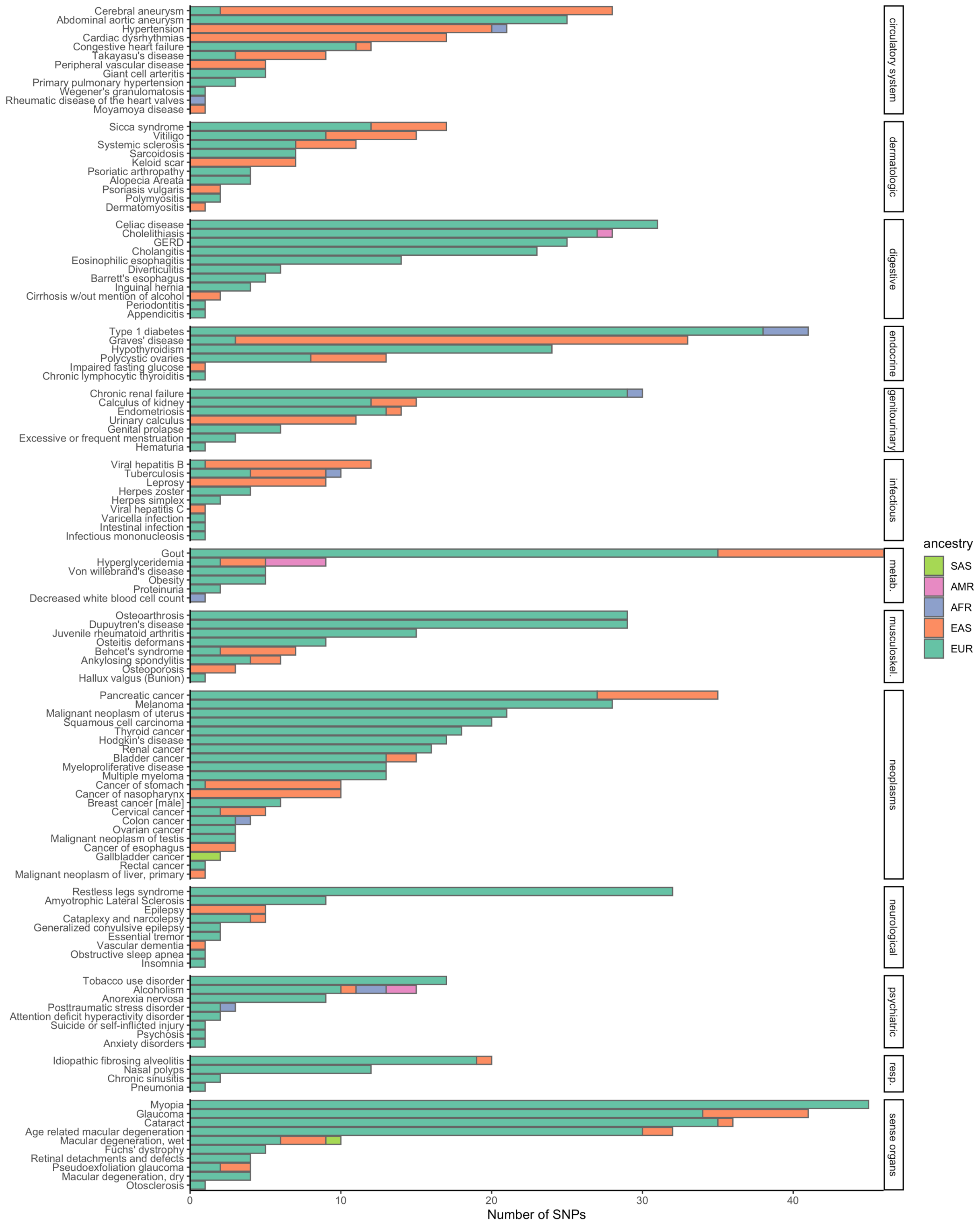

**Figure S2**. Overlap of replications between three European ancestry test cohorts. In total, there were 393 associations that were powered in all three cohorts, 22 (5.6%) of which failed to replicate in all three cohorts.

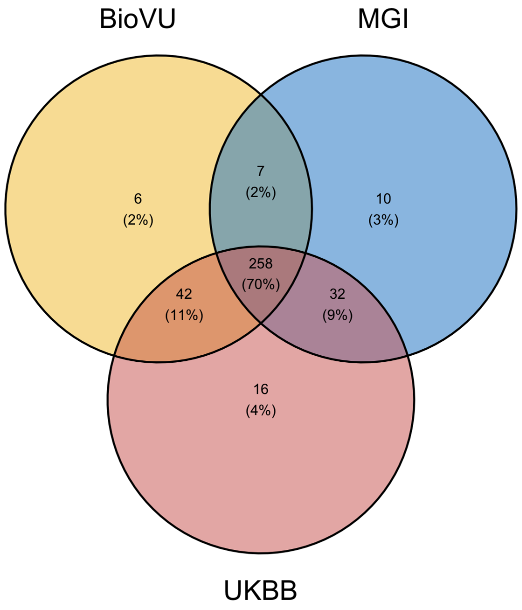

**Figure S3**. Odds ratio comparison between GWAS catalog and test cohorts. Axes were cropped at 3 for visibility.

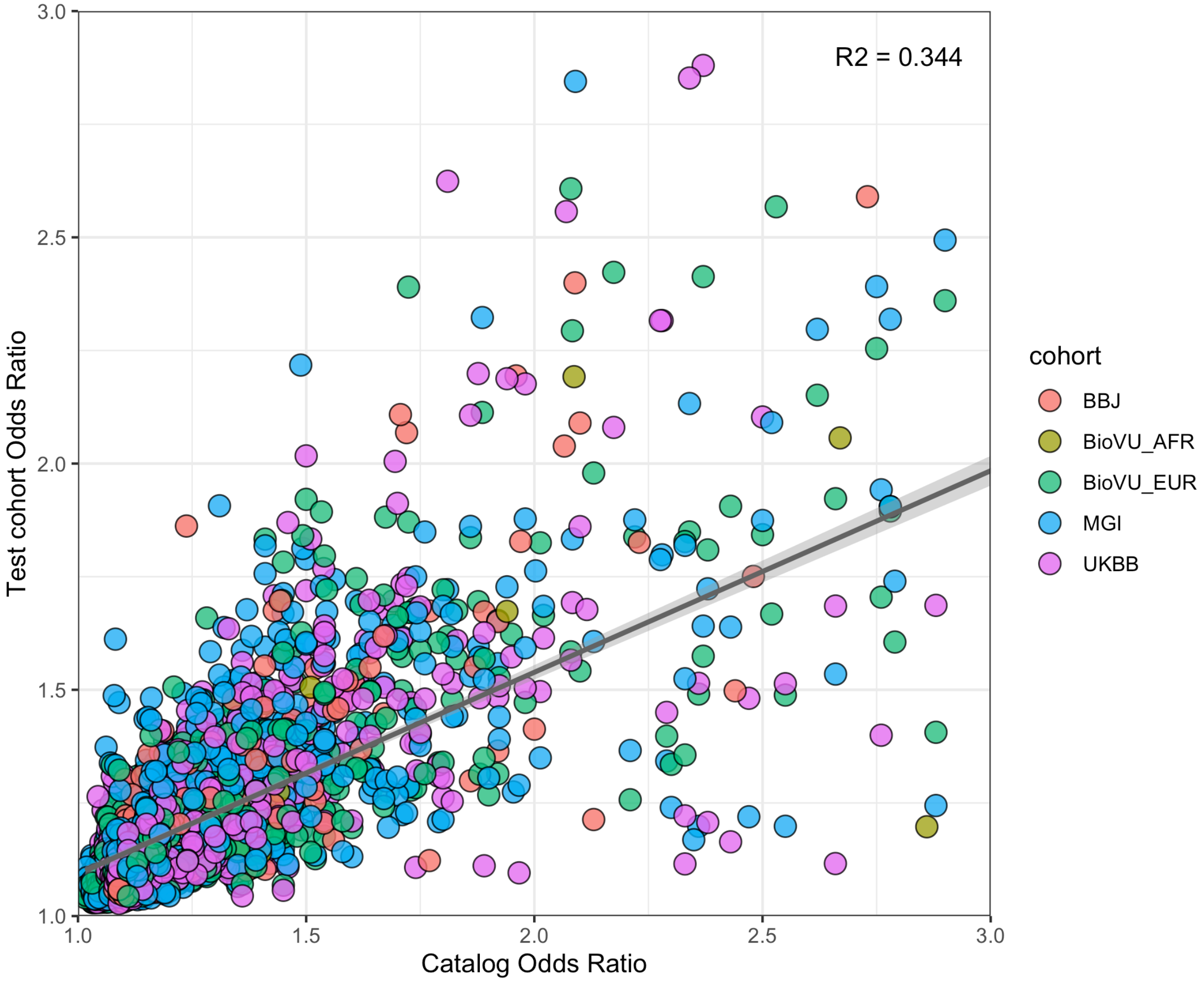
