## Supplement tables 3,4,7,8,9,10 for "The Phenotype-Genotype Reference Map: Improving biobank data science through replication"

**Supplemental Table 3**

| Cohort | # individuals | Phenotype definition | Genotype method | Model | Covariates |
|---|---|---|---|---|---|
| BioVU, Eurpean ancestry | 62,777 | phecodes, MCC=2 with exclude ranges | Illumina MEGA array; Imputed with 1000 Genomes Project Phase 3 | logistic regression implimented in Plink | Sex, age at last visit, top 8 principle components |
| BioVU, African ancestry | 12,142 | phecodes, MCC=2 with exclude ranges | Illumina MEGA array; Imputed with 1000 Genomes Project Phase 3 | logistic regression implimented in Plink | Sex, age at last visit, top 8 principle components |
| MGI | 51,393 | phecodes | Illumina Infinium CoreExome-24 bead array; Imputed with TOPMed | logistic regression implimented in SAIGE | age, inferred sex, genotyping array, and the first ten PCs |
| UKBB | 407,202 | phecodes | UK BiLEVE Axiom array and UK Biobank Axiom array; imputed with TOPMed | logistic regression implimented in SAIGE | age, sex, chip version, and the first four principal |
| BBJ | 178,726 | Custom phecodes based on ICD-10. MCC=1. No exclude ranges | Illumina HumanOmniExpressExome BeadChip and Illumina HumanOmniExpress, HumanExome BeadChip; imputed with 1000 Genomes Project Phase 3 version 5 | generalized linear mixed model implemented in SAIGE (v.0.37) | age, age2, sex, age × sex, age2 × sex and the top 20 principal components |

**Supplemental Table 4**

| phecode | phecode_string | URL |
|---|---|---|
| 010 | Tuberculosis | https://pheweb.jp/pheno/PTB |
| 053 | Herpes zoster | https://pheweb.jp/pheno/Zoster |
| 054 | Herpes simplex | https://pheweb.jp/pheno/Herpes |
| 070.2 | Viral hepatitis B | https://pheweb.jp/pheno/CHB |
| 070.3 | Viral hepatitis C | https://pheweb.jp/pheno/CHC |
| 150 | Cancer of esophagus | https://pheweb.jp/pheno/EsC |
| 151 | Cancer of stomach | https://pheweb.jp/pheno/GaC |
| 153 | Colorectal cancer | https://pheweb.jp/pheno/CRC |
| 155.1 | Malignant neoplasm of liver, primary | https://pheweb.jp/pheno/HepC |
| 157 | Pancreatic cancer | https://pheweb.jp/pheno/PaC |
| 159.3 | Malignant neoplasm of gallbladder and extrahepatic bile ducts | https://pheweb.jp/pheno/BtC |
| 165.1 | Cancer of bronchus; lung | https://pheweb.jp/pheno/LuC |
| 174.11 | Malignant neoplasm of female breast | https://pheweb.jp/pheno/BrC |
| 180.1 | Cervical cancer | https://pheweb.jp/pheno/CeC |
| 184.11 | Malignant neoplasm of ovary | https://pheweb.jp/pheno/OvC |
| 185 | Cancer of prostate | https://pheweb.jp/pheno/PrC |
| 193 | Thyroid cancer | https://pheweb.jp/pheno/ThC |
| 218.1 | Uterine leiomyoma | https://pheweb.jp/pheno/UF |
| 242.1 | Graves' disease | https://pheweb.jp/pheno/GD |
| 244 | Hypothyroidism | https://pheweb.jp/pheno/Hypothyroidism |
| 245.21 | Chronic lymphocytic thyroiditis | https://pheweb.jp/pheno/Hashimoto_Disease |
| 250.1 | Type 1 diabetes | https://pheweb.jp/pheno/T1D |
| 250.2 | Type 2 diabetes | https://pheweb.jp/pheno/T2D |
| 295.1 | Schizophrenia | https://pheweb.jp/pheno/Schizophrenia |
| 296.22 | Major depressive disorder | https://pheweb.jp/pheno/Depression |
| 318 | Tobacco use disorder | https://pheweb.jp/pheno/Smoking_Ever_Never |
| 327.32 | Obstructive sleep apnea | https://pheweb.jp/pheno/SAS |
| 327.4 | Insomnia | https://pheweb.jp/pheno/Insomnia |
| 332 | Parkinson's disease | https://pheweb.jp/pheno/Parkinsons_Disease |
| 345.1 | Epilepsy | https://pheweb.jp/pheno/Epilepsy |
| 361 | Retinal detachments and defects | https://pheweb.jp/pheno/Retinal_Detachment |
| 365 | Glaucoma | https://pheweb.jp/pheno/Glaucoma |
| 366 | Cataract | https://pheweb.jp/pheno/Cataract |

| | | |
|---|---|---|
| 394 | Rheumatic disease of the heart valves | https://pheweb.jp/pheno/Cardiac_Valvular_Disease |
| 411.2 | Myocardial infarction | https://pheweb.jp/pheno/MI |
| 428 | Congestive heart failure; nonhypertensive | https://pheweb.jp/pheno/CHF |
| 433.5 | Cerebral aneurysm | https://pheweb.jp/pheno/CeAn |
| 442.11 | Abdominal aortic aneurysm | https://pheweb.jp/pheno/AA |
| 443.9 | Peripheral vascular disease, unspecified | https://pheweb.jp/pheno/PAD |
| 471 | Nasal polyps | https://pheweb.jp/pheno/Nasal_polyp |
| 475 | Chronic sinusitis | https://pheweb.jp/pheno/Chronic_Sinusitis |
| 476 | Allergic rhinitis | https://pheweb.jp/pheno/Allergic_Rhinitis |
| 480 | Pneumonia | https://pheweb.jp/pheno/Pneumonia |
| 495 | Asthma | https://pheweb.jp/pheno/Asthma |
| 530.11 | GERD | https://pheweb.jp/pheno/GERD |
| 555.2 | Ulcerative colitis | https://pheweb.jp/pheno/UC |
| 571.51 | Cirrhosis of liver without mention of alcohol | https://pheweb.jp/pheno/Cirrhosis |
| 574.1 | Cholelithiasis | https://pheweb.jp/pheno/Cholelithiasis |
| 585.3 | Chronic renal failure [CKD] | https://pheweb.jp/pheno/Chronic_Renal_Failure |
| 615 | Endometriosis | https://pheweb.jp/pheno/Endometriosis |
| 695.42 | Systemic lupus erythematosus | https://pheweb.jp/pheno/SLE |
| 696.41 | Psoriasis vulgaris | https://pheweb.jp/pheno/PsV |
| 697 | Sarcoidosis | https://pheweb.jp/pheno/Sarcoidosis |
| 701.4 | Keloid scar | https://pheweb.jp/pheno/Keloid |
| 709.2 | Sicca syndrome | https://pheweb.jp/pheno/Sjogren_Syndrome |
| 709.4 | Polymyositis | https://pheweb.jp/pheno/PM |
| 711.3 | Behcet's syndrome | https://pheweb.jp/pheno/Behcet_disease |
| 714.1 | Rheumatoid arthritis | https://pheweb.jp/pheno/RA |
| 714.2 | Juvenile rheumatoid arthritis | https://pheweb.jp/pheno/JRA |

### Supplemental Table 7

| | | | | | | | Comparison with previous set (10% less randomization) | | | | | | | | | |
|---|---|---|---|---|---|---|---|---|---|---|---|---|---|---|---|---|
| | | | | | | | | RR, all (Fisher's exact) | | | RR, powered (Fisher's exact) | | | AER, (logistic regression: replication ~ dataset + Powered) | | |
| % random | Assoc. tested | Powered assoc | replicated assoc. | RR, all | RR, powered | AER | Label | OR | CI | P | OR | CI | P | OR | CI | P |
| 0% | 3268 | 853 | 1354 | 41% | 76% | 0.808 | - | | - | - | - | - | - | - | - | - |
| 10% | 3268 | 853 | 1207 | 37% | 71% | 0.720 | 10% vs 0% randomized | 0.83 | (0.7485 to 0.9156) | 2.2E-04 | 0.77 | (0.6197 to 0.9675) | 0.024 | 0.78 | (0.6919 to 0.8713) | 1.7E-05 |
| 20% | 3268 | 853 | 1031 | 32% | 64% | 0.615 | 20% vs 10% randomized | 0.79 | (0.7094 to 0.873) | 5.0E-06 | 0.71 | (0.575 to 0.8738) | 1.1E-03 | 0.73 | (0.6499 to 0.8219) | 1.7E-07 |
| 30% | 3268 | 853 | 942 | 29% | 62% | 0.562 | 30% vs 20% randomized | 0.88 | (0.7894 to 0.9781) | 0.018 | 0.92 | (0.7506 to 1.123) | 0.423 | 0.85 | (0.7522 to 0.9553) | 6.8E-03 |
| 40% | 3268 | 853 | 780 | 24% | 51% | 0.465 | 40% vs 30% randomized | 0.77 | (0.692 to 0.8658) | 6.1E-06 | 0.64 | (0.5283 to 0.784) | 8.6E-06 | 0.73 | (0.6425 to 0.8224) | 4.1E-07 |
| 50% | 3268 | 853 | 579 | 18% | 40% | 0.345 | 50% vs 40% randomized | 0.69 | (0.6076 to 0.7762) | 1.0E-09 | 0.63 | (0.5182 to 0.7678) | 3.0E-06 | 0.64 | (0.5615 to 0.7304) | 3.1E-11 |
| 60% | 3268 | 853 | 780 | 24% | 51% | 0.304 | 60% vs 50% randomized | 0.86 | (0.7521 to 0.9804) | 2.4E-02 | 0.75 | (0.608 to 0.9124) | 4.1E-03 | 0.84 | (0.7338 to 0.9675) | 0.015 |
| 70% | 3268 | 853 | 341 | 10% | 21% | 0.203 | 70% vs 60% randomized | 0.63 | (0.5424 to 0.7313) | 5.9E-10 | 0.55 | (0.4382 to 0.6853) | 6.3E-08 | 0.61 | (0.5215 to 0.7073) | 1.4E-10 |
| 80% | 3268 | 853 | 203 | 6% | 13% | 0.121 | 80% vs 70% randomized | 0.57 | (0.4719 to 0.6836) | 7.1E-10 | 0.55 | (0.4201 to 0.7174) | 6.1E-06 | 0.55 | (0.461 to 0.6672) | 4.1E-10 |
| 90% | 3268 | 853 | 159 | 5% | 6% | 0.095 | 90% vs 80% randomized | 0.77 | (0.6196 to 0.961) | 0.020 | 0.42 | (0.2905 to 0.6026) | 7.4E-07 | 0.77 | (0.6214 to 0.9544) | 0.017 |
| 100% | 3268 | 853 | 79 | 2% | 2% | 0.047 | 100% vs 90% randomized | 0.48 | (0.3634 to 0.6415) | 1.4E-07 | 0.27 | (0.1358 to 0.4974) | 4.8E-06 | 0.48 | (0.3681 to 0.6372) | 2.2E-07 |

**Supplemental table 8**

***Table S8A - exclude ranges analysis***

| Phenotype method | Assoc. tested | Powered assoc. | replicated assoc. | RR, all | % Powered | RR, powered | AER | Comparison | RR, all (Fisher's exact) OR | CI | P | % powered OR | CI | P | RR, powered (Fisher's exact) OR | CI | P | AER, (logistic regression: replication ~ dataset + Powered) OR | CI | P |
|---|---|---|---|---|---|---|---|---|---|---|---|---|---|---|---|---|---|---|---|---|
| With exclude ranges | 3268 | 853 | 1354 | 41.4% | 26.1% | 76.3% | 0.808 | With exclude ranges vs without | 1.03 | (0.93 to 1.14) | 0.580 | 1.07 | (0.8614 to 1.3396) | 0.525 | 1.07 | (0.86 to 1.34) | 0.536 | 1.06 | (0.94 to 1.19) | 0.332 |
| Without exclude ranges | 3268 | 860 | 1331 | 40.7% | 26.3% | 75.0% | 0.788 | | | | | | | | | | | | | |

***Table S8B - Inpatient only analysis***

| Phenotype method | Assoc. tested | Powered assoc. | Replicated assoc. | RR, all | % Powered | RR, powered | AER | Comparison | RR, all (Fisher's exact) OR | CI | P | % powered OR | CI | P | RR, powered (Fisher's exact) OR | CI | P | AER, (logistic regression: replication ~ dataset + Powered) OR | CI | P |
|---|---|---|---|---|---|---|---|---|---|---|---|---|---|---|---|---|---|---|---|---|
| With all codes | 3268 | 853 | 1354 | 41.4% | 26.1% | 76.3% | 0.808 | With all codes vs inpatient only | 1.69 | (1.5165 to 1.8835) | 4.0E-22 | 2.28 | (1.9873 to 2.6082) | 3.3E-35 | 1.01 | (0.7603 to 1.3433) | 0.942 | 0.56 | 1.0387 | (0.9155 to 1.1784) |
| With inpatient only codes | 2806 | 377 | 828 | 29.5% | 13.4% | 76.1% | 0.814 | | | | | | | | | | | | | |

**Supplemental table 9**

| | | | | | | | | Comparison with previous set (10% less randomization) | | | | | | | | | | | | |
|---|---|---|---|---|---|---|---|---|---|---|---|---|---|---|---|---|---|---|---|---|
| | | | | | | | | | RR, all (Fisher's exact) | | | % Powered (Fisher's exact) | | | RR, powered (Fisher's exact) | | | AER, (logistic regression: replication ~ dataset + Powered) | | |
| **MCC** | **associations tested** | **RR (all)** | **# replicated** | **# powered** | **% powered** | **RR (powered)** | **AER** | **Label** | **OR** | **CI** | **P** | **OR** | **CI** | **P** | **OR** | **CI** | **P** | **OR** | **CI** | **P** |
| 1 | 3,296 | 39.5% | 1,303 | 1,126 | 34.2% | 67.5% | 0.671 | - | - | - | - | - | - | - | - | - | - | - | - | - |
| 2 | 3,268 | 41.4% | 1,354 | 853 | 26.1% | 76.3% | 0.808 | MCC 2 vs 1 | 1.08 | (0.9792 to 1.1956) | 0.119 | 0.68 | (0.6112 to 0.758) | **1.2E-12** | 1.55 | (1.2634 to 1.9089) | **1.9E-05** | 1.50 | (1.3332 to 1.6765) | **6.0E-12** |
| 3 | 3,242 | 39.7% | 1,287 | 753 | 23.2% | 77.6% | 0.838 | MCC 3 vs 2 | 0.93 | (0.8419 to 1.0287) | 0.158 | 0.86 | (0.7638 to 0.9604) | **7.5E-03** | 1.07 | (0.8439 to 1.3633) | 0.593 | 1.07 | (0.956 to 1.2045) | 0.232 |
| 4 | 3,224 | 37.6% | 1,211 | 666 | 20.7% | 79.1% | 0.842 | MCC 4 vs 3 | 0.91 | (0.8257 to 1.0114) | 0.078 | 0.86 | (0.7634 to 0.97) | **0.013** | 1.10 | (0.8449 to 1.426) | 0.479 | 1.00 | (0.8908 to 1.1268) | 0.975 |
| 5 | 3,174 | 36.8% | 1,167 | 602 | 19.0% | 81.1% | 0.865 | MCC 5 vs 4 | 0.97 | (0.8722 to 1.0711) | 0.518 | 0.90 | (0.7933 to 1.0186) | 0.090 | 1.13 | (0.8484 to 1.5043) | 0.399 | 1.05 | (0.9309 to 1.1823) | 0.432 |
| 6 | 3,141 | 35.8% | 1,126 | 534 | 17.0% | 81.3% | 0.883 | MCC 6 vs 5 | 0.96 | (0.8662 to 1.0663) | 0.448 | 0.88 | (0.7678 to 0.9973) | 0.042 | 1.01 | (0.7444 to 1.3822) | 0.940 | 1.04 | (0.9179 to 1.1696) | 0.566 |
| 7 | 3,095 | 35.6% | 1,102 | 496 | 16.0% | 81.3% | 0.903 | MCC 7 vs 6 | 0.99 | (0.8909 to 1.099) | 0.853 | 0.93 | (0.8131 to 1.0675) | 0.306 | 1.00 | (0.7211 to 1.3834) | 1.000 | 1.04 | (0.9226 to 1.1777) | 0.505 |
| 8 | 2,998 | 36.3% | 1,087 | 458 | 15.3% | 82.8% | 0.932 | MCC 8 vs 7 | 1.03 | (0.9252 to 1.1438) | 0.612 | 0.94 | (0.8207 to 1.0876) | 0.438 | 1.11 | (0.7846 to 1.5646) | 0.556 | 1.07 | (0.9455 to 1.2096) | 0.285 |

**Supplemental table 10**

*Associations that did not replicate in UKB, MGI, or BioVU*

| assoc_ID | rsID | phecode | phecode_string | category_string | Study_accession |
|---|---|---|---|---|---|
| 99743 | rs35509282 | 153 | Colorectal cancer | neoplasms | GCST002528 |
| 189758 | rs7202877 | 250.1 | Type 1 diabetes | endocrine | GCST000392 |
| 81628 | rs1424233 | 278.1 | Obesity | metabolic/heme | GCST000317 |
| 294914 | rs2116830 | 278.1 | Obesity | metabolic/heme | GCST001130 |
| 37646 | rs17616243 | 295.1 | Schizophrenia | psychiatric | GCST007219 |
| 296648 | rs7527939 | 295.1 | Schizophrenia | psychiatric | GCST001733 |
| 80352 | rs1012053 | 296.1 | Bipolar | psychiatric | GCST000033 |
| 312906 | rs1709393 | 300 | Anxiety disorders | psychiatric | GCST003370 |
| 188159 | rs7590720 | 317.1 | Alcoholism | psychiatric | GCST000432 |
| 188443 | rs1835740 | 340 | Migraine | neurological | GCST000782 |
| 193678 | rs284489 | 365.1 | Open-angle glaucoma | sense organs | GCST001493 |
| 23881 | rs6941513 | 411.2 | Myocardial infarction | circulatory system | GCST003430 |
| 273727 | rs6922269 | 411.4 | Coronary atherosclerosis | circulatory system | GCST000057 |
| 19376 | rs3131623 | 480 | Pneumonia | respiratory | GCST005009 |
| 36977 | rs75801644 | 615 | Endometriosis | genitourinary | GCST004873 |
| 36998 | rs644045 | 615 | Endometriosis | genitourinary | GCST004873 |
| 37009 | rs10129516 | 615 | Endometriosis | genitourinary | GCST004873 |
| 204280 | rs11073328 | 695.42 | Systemic lupus erythematosus | dermatologic | GCST002463 |
| 204281 | rs8023715 | 695.42 | Systemic lupus erythematosus | dermatologic | GCST002463 |
| 81745 | rs13017599 | 714.1 | Rheumatoid arthritis | musculoskeletal | GCST000420 |
| 272954 | rs3761847 | 714.1 | Rheumatoid arthritis | musculoskeletal | GCST000070 |
| 194828 | rs11842874 | 740 | Osteoarthrosis | musculoskeletal | GCST001209 |
